# phyddle: software for exploring phylogenetic models with deep learning

**DOI:** 10.1101/2024.08.06.606717

**Authors:** Michael J. Landis, Ammon Thompson

## Abstract

Phylogenies contain a wealth of information about the evolutionary history and process that gave rise to the diversity of life. This information can be extracted by fitting phylogenetic models to trees. However, many realistic phylogenetic models lack tractable likelihood functions, prohibiting their use with standard inference methods. We present `phyddle`, pipeline-based software for performing phylogenetic modeling tasks on trees using likelihood-free deep learning approaches. `phyddle` has a flexible command-line interface, making it easy to integrate deep learning approaches for phylogenetics into research workflows. `phyddle` coordinates modeling tasks through five pipeline analysis steps (*Simulate, Format, Train, Estimate*, and *Plot*) that transform raw phylogenetic datasets as input into numerical and visual model-based output. We conduct three experiments to compare the accuracy of likelihood-based inferences against deep learning-based inferences obtained through `phyddle`. Benchmarks show that `phyddle` accurately performs the inference tasks for which it was designed, such as estimating macroevolutionary parameters, selecting among continuous trait evolution models, and passing coverage tests for epidemiological models, even for models that lack tractable likelihoods. Learn more about `phyddle` at https://phyddle.org.

## Motivation

Good phylogenetic model design balances biological, statistical, and computational considerations (Felsenstein 1985; Hansen and Martins 1996; Kelchner and Thomas 2007; Rodrigue and Philippe 2010; Servedio et al. 2014; Rolland et al. 2023). Not only should a phylogenetic model realistically describe how evolutionary lineages change across generations, it should allow accurate estimates to be obtained from finite data in a short amount of time. In practice, phylogenetics models are often fitted to data through the likelihood function, e.g. using maximum likelihood or Bayesian approaches. Likelihood is central to statistical modeling in general, with phylogenetic modeling being no exception.

It is not overstated to say likelihood-based inference revolutionized phylogenetics (Cavalli-Sforza and Edwards 1967; Felsenstein 1973).

That said, phylogenetic models with desirable biological or statistical properties may not yield tractable likelihood functions. In such cases, researchers face a dilemma: either use tractable but suboptimal phylogenetic models, or invest substantial resources and effort to design and validate sophisticated likelihood-based inference methods for more-realistic models. Inference methods that do not require the true model likelihood (e.g. Beaumont et al. 2002; Wood 2010; Rue et al. 2009) provide a third option. Among these methods, simulation-based deep learning approaches have proven useful for many general model estimation tasks (Tejero-Cantero et al. 2020; Sainsbury-Dale et al. 2024;

Lenzi et al. 2023; Rmus et al. 2024). Moreover, newer deep learning approaches have made progress on a variety of foundational modeling problems in phylogenetics (Borowiec et al. 2022; Mo et al. 2024), including parameter estimation (Voznica et al. 2022; Lambert et al. 2023; Thompson et al. 2024), model selection (Voznica et al. 2022), and tree inference (Suvorov et al. 2020; Kong et al. 2023; Nesterenko et al. 2024; Smith and Hahn 2023).

This application note introduces `phyddle`, a phylogenetic modeling framework for training neural networks with simulated data through supervised learning (LeCun et al. 2015; Goodfellow et al. 2016). A primary goal of `phyddle` is to equip biologists with a deep learning pipeline workflow so they may explore and apply realistic, but otherwise intractable, phylogenetic models in empirical settings. To do so, `phyddle` automates and streamlines many routine aspects of phylogenetic deep learning, allowing biologists to instead focus on data analysis tasks, such as modeling, simulation, validation, and interpretation. `phyddle` generalizes existing deep learning techniques (Bokma 2006; Voznica et al. 2022; Lambert et al. 2023; Thompson et al. 2024) for modeling lineage diversification (e.g. Maddison et al. 2007; Stadler and Bonhoeffer 2013) and trait evolution (e.g. Pagel 1994; Hansen and Martins 1996) to a broader class of user-defined phylogenetic models and scenarios. Currently supported modeling tasks include parameter estimation, model selection, the inference phylodynamic trends, and ancestral root state reconstruction from fixed phylogenies and character data as input. Future versions of `phyddle` will support other classical modeling tasks in phylogenetics, such as divergence time estimation and tree inference.

The remainder of this overview assumes some basic familiarity with deep learning concepts and practices (see Borowiec et al. 2022 or Goodfellow et al. 2016 for background reading). We highlight only some of `phyddle`’s features below, and refer readers to the documentation for technical descriptions of current software functionality: https://phyddle.org.

## Design & Features

`phyddle` uses deep learning and simulation-trained neural networks to perform phylogenetic modeling tasks. A standard `phyddle` analysis is executed through a pipeline built from five modular steps: *Simulate, Format, Train, Estimate*, and *Plot* (Figure 1).

**Figure 1:**
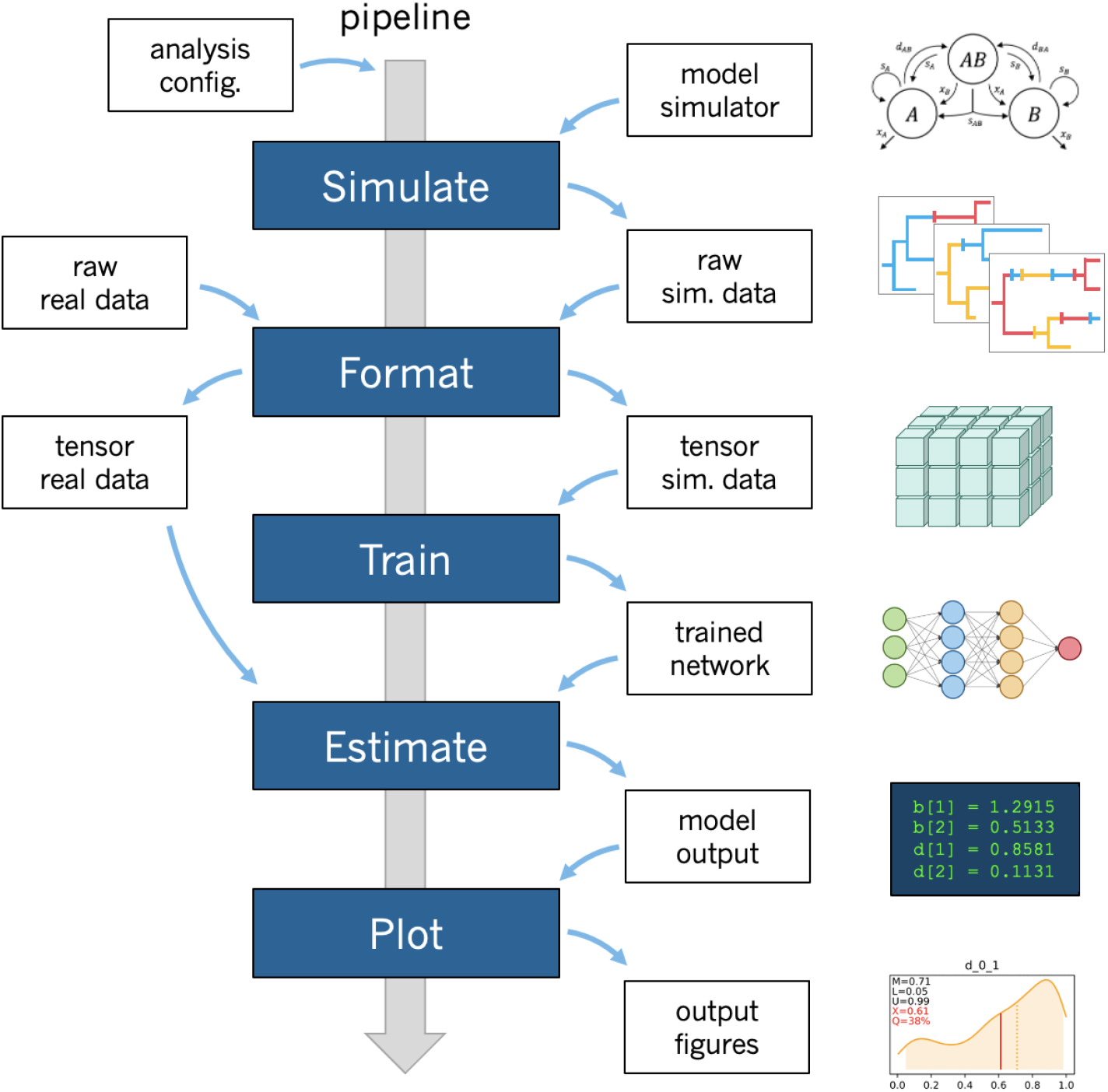
Overview of `phyddle` workflow. Researchers provide the software with a configuration file and a simulation script that determine the analysis settings. Upon running `phyddle`, the *Simulate* step produces a raw (unformatted) pool of training datasets that represent data-generating parameters (labels) and realizations of the data-generating model (examples). Next, the *Format* step restructures the raw training data into tensors, while also reclassifying simulating conditions (e.g. parameters) as either data (treated as observed) or labels (to be predicted). The *Train* step loads the tensor data, constructs and fits neural network architecture, saves a copy of the trained network, and reports training results. The *Estimate* step generates model predictions for test and empirical datasets that were not used during training. Lastly, the *Plot* step produces figures and reports that summarize the pipeline results.

Settings for each step are controlled through command-line options and customizable configuration files. Being a pipeline, `phyddle` output is stored in a predictable manner, allowing the output from one step to become the input for downstream steps. `phyddle` functions as both an interactive command-line tool and as a scriptable interface, allowing researchers to design and debug workflows line-by-line before writing scripts to automate repetitive and large-scale jobs.

The full pipeline begins with the *Simulate* step. This step generates large numbers of example datasets under a model with a user-specified simulation script. The *Simulate* step is intended to be as flexible and simple as possible so researchers may focus on model design. Users provide `phyddle` with model simulators written in whichever programming language suits their needs. Each simulation script itself defines the phylogenetic model, simulates data under that model, and saves all relevant information using standard file formats required by `phyddle`. When run, each simulation generates a training example, comprised of data (e.g. a phylogenetic tree and a character matrix) and labels (e.g. the data-generating parameters). Simulators can also record statistics of phylogenetic and character data patterns that aid inference (e.g. summary statistics Saulnier et al. 2017; Janzen and Etienne 2024) or that act as prediction targets (e.g. ancestral root states, ancestral lineage counts). Simulation tasks may be run in batches and across processors, in parallel. The full set of gathered simulations comprise the raw and unformatted training dataset. Users are encouraged to either modify the simulator scripts bundled with `phyddle`, written using R (R Core Team 2013), Python (Van Rossum and Drake 2009), RevBayes (HÖhna et al. 2016), MASTER (Vaughan and Drummond 2013), and PhyloJunction (Mendes and Landis 2024), or to write their own script from scratch using a language of their choice.

Next, the *Format* step encodes raw datasets into a set of tensors, analogous to *N* -dimensional arrays of variables, which are later processed by the *Train* step. *Format* processes all valid examples from *Simulate*, plus any empirical datasets flagged for analysis by `phyddle`. *Format* converts each raw dataset into three types of tensors: a phylogenetic data tensor, an auxiliary data tensor, and a label tensor.

Phylogenetic data tensors encode phylogenetic and tip-state data into a 2D tensor structure suitable for supervised learning with convolutional neural network (CNN) layers.

`phyddle` encodes phylogenetic data with the compact bijective ladderized vector (CBLV; Voznica et al. 2022) for serially sampled trees and the compact diversity vector (CDV; Lambert et al. 2023) for extant-only trees, with options to include additional rows of branch length information, such as branch lengths and node heights. `phyddle` also allows users to associate multiple categorical or numerical traits with each taxon (+S; Thompson et al. 2024). These tree encodings have small memory/storage footprints that are linearly proportional to taxon count and standardize valuable node-order and branch-length information contained within the input trees (Voznica et al. 2022; Lambert et al. 2023).

Auxiliary data tensors are used as input for dense feed-forward neural network (FFNN) layers. Each row in the auxiliary data tensor represents one raw dataset, with each column corresponding to a “known” model parameter for the data-generating process (e.g. population size, sampling effort) or a summary statistic of the data pattern (e.g. tree height, tree balance, etc.). During *Format*, `phyddle` includes its own standard summary statistics, such as the number of taxa, number of characters, state frequencies, branch length statistics, tree balance statistics, and more.

Label tensors define the prediction targets (outputs). Each row in the label tensor references a single, raw dataset with columns corresponding to different model parameters or other prediction targets. Labels can have numerical (e.g. rates, species counts) or categorical values (e.g. model type, ancestral states).

Once formatted into tensors, *Format* subdivides the simulated dataset into a training dataset, to optimize the network during *Train*, and a test dataset, to later assess trained network performance during *Estimate*. Training, test, and empirical tensors are then saved separately, either in human-readable .csv format or in a compact .hdf5 format that is often 20x smaller than its .csv counterpart.

The *Train* step constructs, trains, calibrates, and saves a neural network and its training history. Neural networks are built and trained using PyTorch (Paszke et al. 2019) with the option to use CPU and Nvidia GPU (CUDA) parallelization. `phyddle` combines convolutional and pooling layers used in convolutional neural networks (CNN) and fully-connected, dense layers used in feed-forward neural networks (FFNN) for its network architecture (Goodfellow et al. 2016; Borowiec et al. 2022). Phylogenetic data tensors are processed as input using a set of CNN layers, whereas the auxiliary data tensors are processed with FFNN layers. The output of these CNN and FFNN layers are then concatenated and passed into a final series of dense FFNN layers for each training target that terminate with an output layer for prediction.

Once the network is constructed, `phyddle` uses supervised learning with training examples from *Simulate* to train, validate, and calibrate prediction intervals from the neural network. Networks are trained by minimizing a loss function, which measures error between network predictions and true label values. Training ends after a fixed number of epochs or when an early-stopping condition is met (e.g. loss scores for validation examples worsen as training progresses). `phyddle` networks can be trained to estimate categorical labels and numerical labels. After training, `phyddle` uses previous point estimates and new calibration examples to produce conformalized prediction intervals (CPIs) with coverage guarantees (Romano et al. 2019). For example, 80% CPIs for new predictions will contain the true parameter value 80% of the time, assuming the training and new data are generated by the same model. `phyddle` allows users to adjust important aspects of training and network architecture through the configuration file, including: depths and widths of layers, convolutional kernel settings (size, stride, dilation, etc.), activation functions, loss functions, the optimizer, and proportions of simulated data assigned to training, test, validation, and calibration example datasets.

When complete, the trained network is saved to file in .hdf5 format so it may be used for future estimation tasks. The history of network performance (e.g. loss and accuracy) for the training and validation datasets is recorded in a .csv file to identify training failures. Tables of predicted versus true values for the training and test datasets are also saved in .csv files. Lastly, `phyddle` stores the means and standard deviations needed to standardize new biological datasets for use with the trained network for predictions.

The *Estimate* step uses the trained network to make predictions from new datasets. When called, *Estimate* generates predictions for the test dataset, which was withheld from training, to assess performance of the trained network. *Estimate* also runs against any empirical datasets identified by the analysis. As a safety feature, this step warns users of empirical samples detected as out-of-distribution (i.e. outliers relative to the training dataset) and that corresponding predictions may not be trustworthy. *Estimate* is the simplest and fastest step, generally producing new estimates from the trained network within milliseconds. After the initial cost of simulating data and training and validating the network, new estimates from the network are effectively free.

Lastly, the *Plot* step visualizes the output from a `phyddle` analysis, beginning with empirical results and ending with network performance and architecture. Estimates for test and empirical examples for numerical and categorical variables are shown first. Empirical data and estimates are also plotted against the marginal densities and dimension-reduced joint densities of the training data and labels. Prediction accuracy for training and test datasets are displayed as scatter plots (with prediction intervals) for numerical labels and confusion matrices for categorical labels. Training metrics are plotted as loss and accuracy scores across epochs for training and validation datasets. The network architecture is shown last. After reviewing the visualizations and metrics from *Plot*, users can adjust the pipeline settings (e.g. more simulations, deeper network architecture, more training epochs, faster learning rate, etc.) and then rerun the relevant steps to improve network performance.

We note that, as with maximum likelihood and Bayesian approaches, results from deep learning approaches also must be carefully validated before being trusted (Goodfellow et al. 2016; Borowiec et al. 2022). `phyddle` is equipped with automated features to promote the safe use of the software, such as: verification that training examples are correctly formatted and present (*Simulate* and *Format*); warnings when empirical data and/or estimates fall outside the distribution of training examples (*Estimate* and *Plot*)); early-stopping rules (*Train*) and training history visualizations (*Plot*) to prevent network underfitting and overfitting; warnings for poor prediction accuracy and coverage among training and test examples to identify poor network performance (*Train, Estimate*, and *Plot*); and a customizable pipeline design to easily explore the sensitivity of different analysis and network settings upon prediction performance (*Train* and *Plot*). Researchers inexperienced with deep learning are cautioned that modeling errors caused by information leakage from the analyst into the training procedure, the incorrect implementation of a simulator, and the incoherent definition of a model are also concerns, but factors such as these cannot be automatically detected by `phyddle`. Researchers should not treat any method, `phyddle` included, as a “black box” that always generates statistically and biologically meaningful results. Visit https://phyddle.org/overview.html#safe-usage to learn how to use the software safely.

The modular design of the `phyddle` pipeline makes it relatively easy to specify, train, save, and share new models with other biologists. Only a configuration file and the contents of the *Train* directory are needed for other researchers to perform *Estimate* tasks with the trained network on their empirical datasets. Entire project workspaces can be archived to ensure research is reproducible.

To showcase the flexibility and jump-start the adoption of `phyddle`, it is bundled with example configuration files, simulation scripts, and pipeline results for a variety of phylogenetic models (https://github.com/mlandis/phyddle/workspace). Current examples include state-dependent speciation-extinction models (SSE), susceptible-infectious-recovered models (SIR), and trait evolution models that variously use R, Python, RevBayes (HÖhna et al. 2016), PhyloJunction (Mendes and Landis 2024), and MASTER (Vaughan and Drummond 2013) to simulate training datasets. These generic examples are meant to be adapted by biologists developing customized `phyddle` pipelines to model their systems.

## Example analyses

We applied `phyddle` to three phylogenetic modeling scenarios to illustrate how the deep learning software behaves relative to existing likelihood-based methods.

We trained one network per experiment. Each network has two separate input layers and one or more separate output layers. The first input layer processes phylogenetic data tensors by feeding information into three parallel series of 1D convolutional and pooling layers: the first series has three convolutional layers with 64, 96, and 128 output channels and kernel sizes of 3, 5, and 7; the second series has two convolutional layers with 64 and 96 output channels, kernel sizes of 7 and 9, and strides of 3 and 6; and the third series has two convolutional layers with 32 and 64 output channels, kernel sizes of 3 and 5, and dilation sizes of 3 and 5. Each of the three convolutional layer-series feeds into a 1D adaptive average pooling layer. The second input layer processes auxiliary data tensors (summary statistics and “known” parameters) through three dense layers with 128, 64, and 32 nodes, in that order. All four series of layers then unite through a concatenation layer, which feeds into one or more dense layer series, one for each output layer needed by the experiment. Dense layers in each of these series contain 128, 64, and 32 nodes. When trained to predict numerical labels (e.g. rates), the network has three output layers for the point estimate and the lower and upper bound prediction interval estimates. When predicting categorical labels (e.g. model categories, ancestral states), the network has one softmax output layer per label. Rectified linear units were used as activation functions (Nair and Hinton 2010). Loss values were computed as the sum of mean-squared error values for numerical point estimates, pinball loss values for the numerical prediction interval bound estimates, and cross-entropy values for categorical estimates. For training, the full simulated dataset was split into training (70%), validation (5%), test (5%) and calibration (20%) data subsets. Networks were trained using stochastic batch gradient descent and the Adam optimizer (Kingma and Ba 2014) with a learning rate of 0.001 and *β* = (0.9, 0.999). Networks were trained for up to 200 epochs, but stopped early if validation loss scores worsened across three consecutive epochs.

After training, we reviewed the results from the *Plot* step to validate each network was properly trained. For each analysis, we confirmed the total loss scores for the validation and training datasets were similar and that training stopped once the validation loss score increased for three consecutive epochs. We also confirmed that parameter prediction accuracy for train and test parameter estimates were both similar to each other and to the truth, and that predicted coverage was similar to the target coverage level. Lastly, we compared network-based and likelihood-based parameter estimates against each other.

We ran all deep learning analyses through PyTorch (Paszke et al. 2019) using `phyddle` 0.2.2 on a server with 2x Intel CPUs (112 cores total), 1x Nvidia RTX A4000 GPU, and 128GB RAM. We report approximate runtimes for all analyses using this hardware.

Saved workspace projects for the three example analyses are hosted at https://github.com/mlandis/phyddle_ms. These projects include configuration files, simulator scripts, and results visualized by *Plot*.

### Macroevolution under BiSSE

To demonstrate the use of `phyddle` for a parameter estimation task, we simulated phylogenetic datasets under a simple binary state-dependent speciation-extinction (BiSSE; Maddison et al. 2007). The BiSSE model is a tree-generating process that assumes that speciation rates and extinction rates depend on the state of an evolving binary character. Our simulation used the following settings:

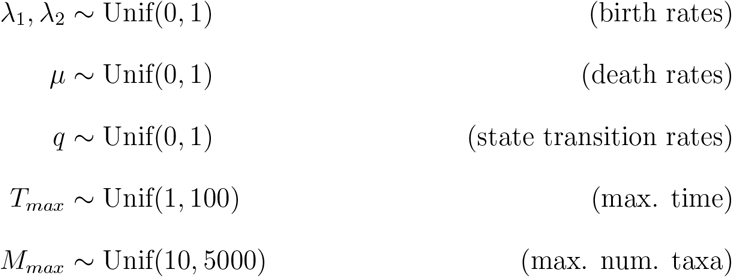

where *λ*_1_ and *λ*_2_ are state-dependent birth rates for lineages in states 1 and 2 respectively, *µ* is the state-independent death rate, and *q* is the symmetric transition rate between states 1 and 2. Each simulation was terminated after *T*_*max*_ units of evolutionary time or when the phylogeny contained *M*_*max*_ extant taxa, whichever condition arose first. Only extant taxa were retained.

We used the R packages `ape` (Paradis and Schliep 2019) and `castor` (Louca and Doebeli 2018) to simulate and fit phylogenies and character matrices with a BiSSE model. We simulated datasets with the `castor::simulate dsse()` function. Datasets with 500 or fewer taxa included all taxa and had sampling fractions of 1, whereas larger trees were downsampled to 500 taxa and recorded as having sampling fractions of less than 1. All data-generating parameters and the sampling fraction were log-transformed before being saved to file for the example dataset. Evolutionary rates were treated as estimation targets, while the sampling fraction was treated as a known parameter. Simulated trees were saved in Newick format, and data matrices were stored as comma-separated value tables. This simulation procedure is stored in the file `sim bisse.R`, as referenced in the `phyddle` configuration file below.

Maximum likelihood estimates (MLEs) for each dataset were obtained using the `castor::fit musse()` function under the `nlminb` optimization method. To improve optimization success, each MLE represents 10 trials with 30 initial guesses (“scouts”) per trial and one “noisy” guess that is centered on the true data-generating parameters. Optimization was further restricted to search for parameter values within the simulated bounds (minimum 0, maximum 1), as it improved MLE parameter accuracy. We also expect MLE might mildly underperform in this analysis, because the inference model conditions on survival of the tree, rather than the *T*_*max*_ and *M*_*max*_ stopping criteria used for the simulating model.

Neural network estimates were generated with `phyddle`, using a config file that is partly shown in Listing 1. As the comments explain, this config expects extant-only trees with one character and two states per taxon. Four numerical parameters (log-transformed *λ*_1_, *λ*_2_, *µ*, and *q*) are estimated, while one numerical parameter (log-transformed sampling proportion, *ρ*) is treated as known, as stated earlier. Default `phyddle` settings are used for any unspecified settings in the config file (see documentation).

**Listing 1:**
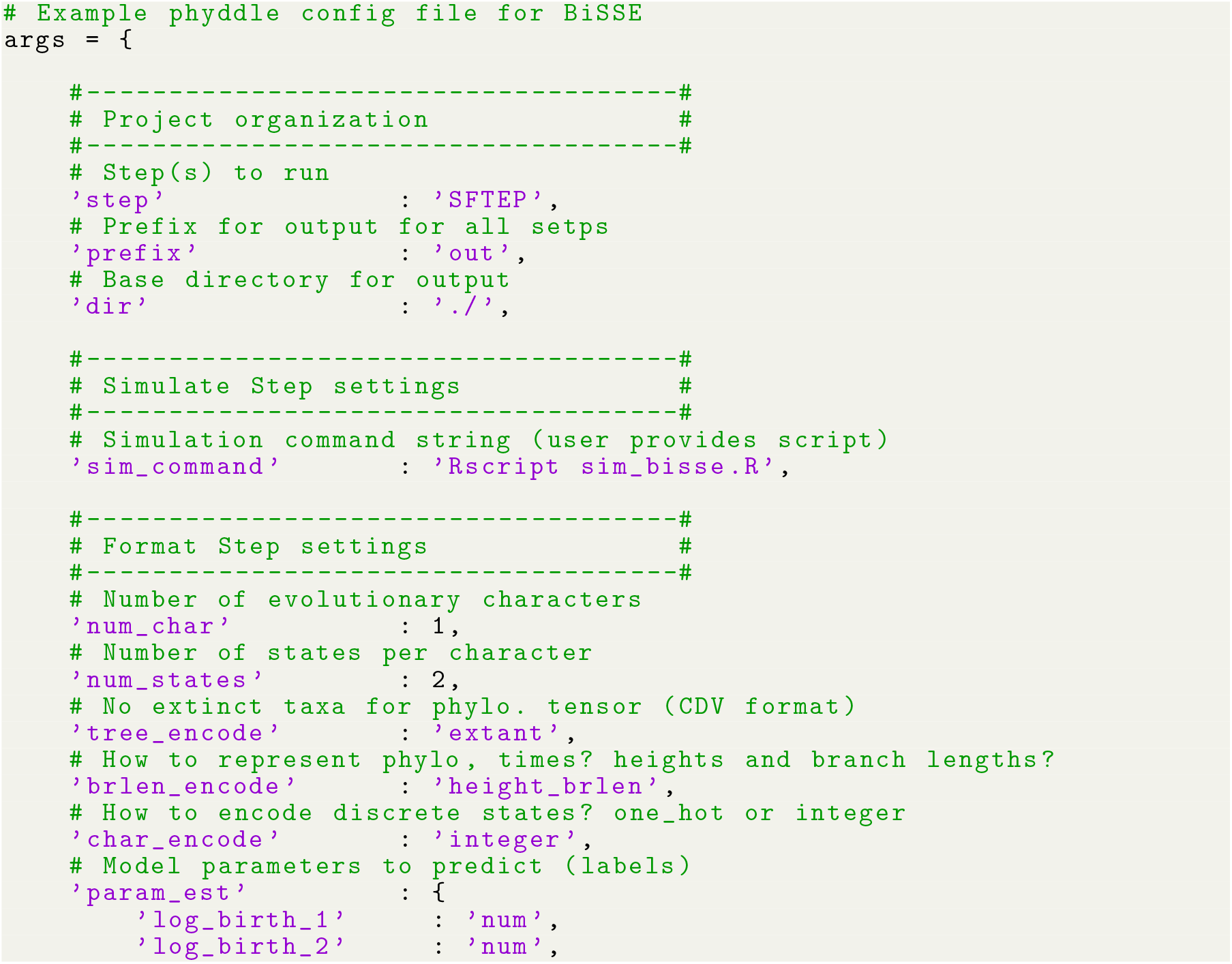
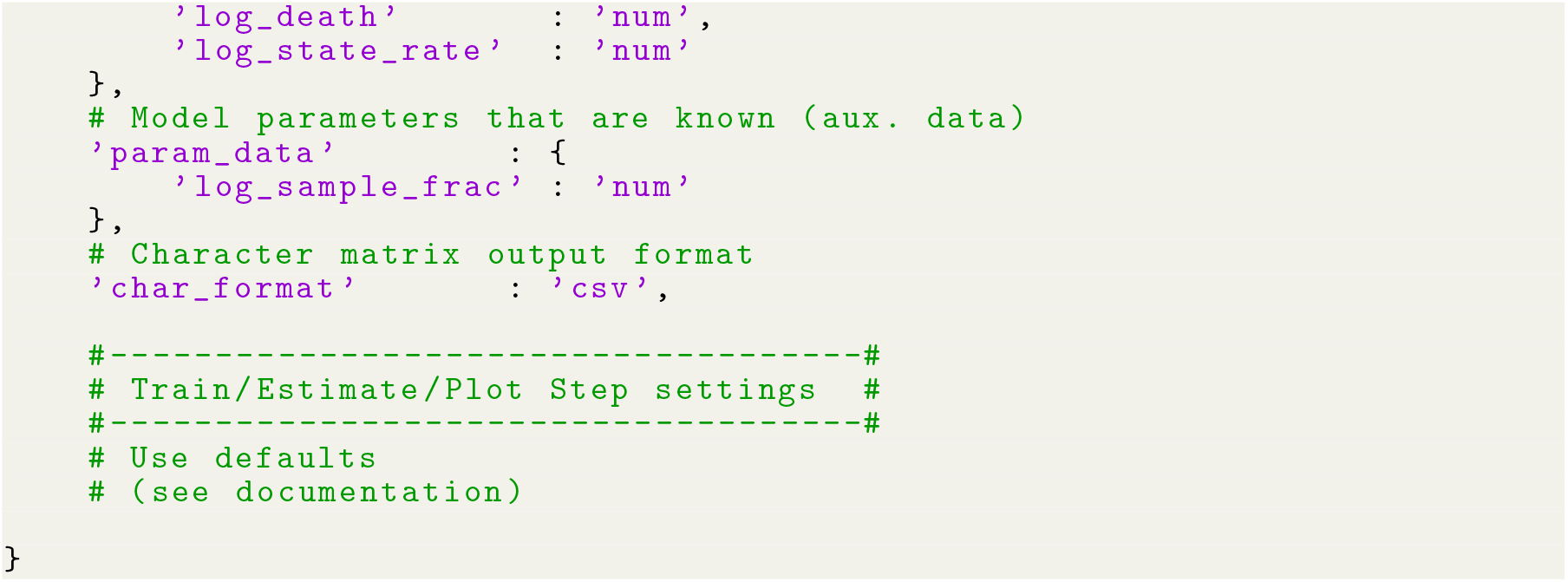
Minimal `phyddle` config file for BiSSE analysis. We specify only a few key settings and let `phyddle` apply default values to the remaining settings (see documentation).

We simulated 50,000 example datasets for the `phyddle` analysis. The configuration file, simulation script, and pipeline results are stored in the project archive, `bisse project.tar.gz`. Pipeline results can be reproduced from scratch with the commands in Listing 2:

**Listing 2:**
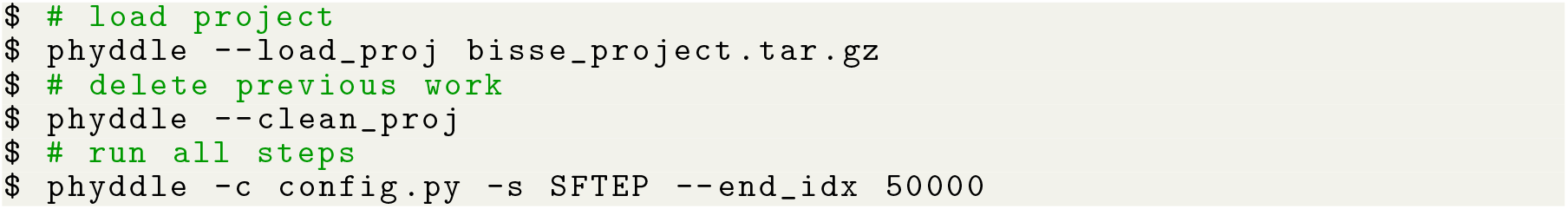
Shell commands for BiSSE analysis using `phyddle`.

We used 100 simulated datasets to compare the performance between `phyddle` and maximum likelihood. Parameter estimates for `phyddle` (Fig. 2A-D) and maximum likelihood (Fig. 2E-H) are each are highly correlated with the true data-generating parameters. In addition, `phyddle` and maximum likelihood methods tend to agree with each other as much as the truth (Fig. 2I-L), implying both methods extract similar information from the same data pattern. `phyddle` required approximately 4m to simulate and format data, 1.5m to train the network, and less than 5s to estimate parameters (5.5m total), whereas our parallelized maximum likelihood script needed about 1.5m.

**Figure 2:**
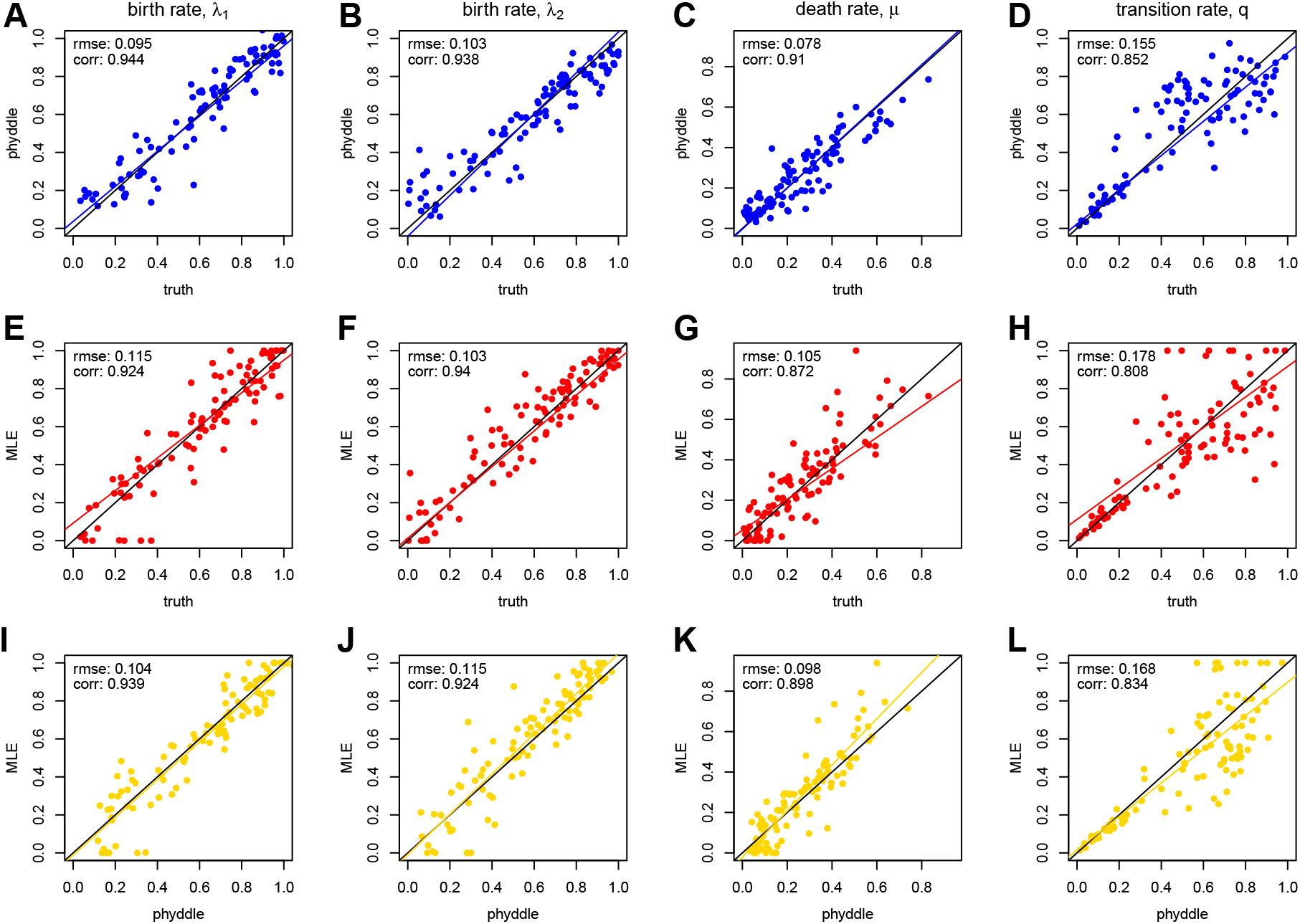
Comparison of maximum likelihood and `phyddle` parameter estimates. Each column corresponds to a BiSSE model rate that was estimated: *λ*_1_, *λ*_2_, *µ*, and *q*. The first row compares `phyddle` estimates to true parameter values (blue, A-D), the second row compares MLEs to true values (red, E-H), and the third row compares `phyddle` and MLE estimates to each other (gold, I-L). The black line is set to intercept-0 and slope-1, representing perfectly matching values. The colored lines are standard linear regressions, and align with the black line when estimates agree with the truth (blue and red) or with each other (gold).

### Continuous trait evolution model selection

We next demonstrate `phyddle`’s ability to predict categorical variables in a model selection task. We simulated datasets under four models that represent alternate modes of continuous trait evolution: Brownian motion for incremental change (BM; Felsenstein 1985), Ornstein-Uhlenbeck for stationarity (OU; Hansen and Martins 1996), Early Burst for explosive change (EB; Harmon et al. 2010), and a Normal Inverse Gaussian Lévy process for pulsed change (LP; Landis and Schraiber 2017). Trees were simulated with a standard constant-rate birth-death model, under the following conditions

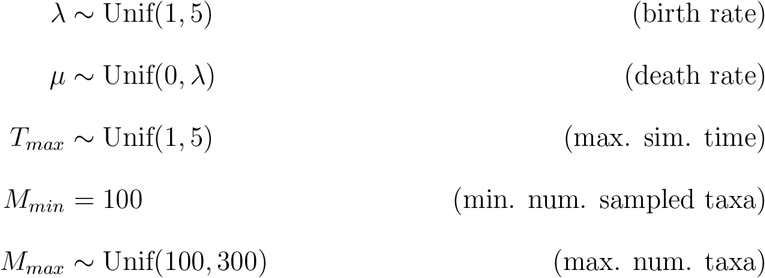

Next, for each tree, we first sampled a model type and its corresponding model parameters and then simulating the dataset:

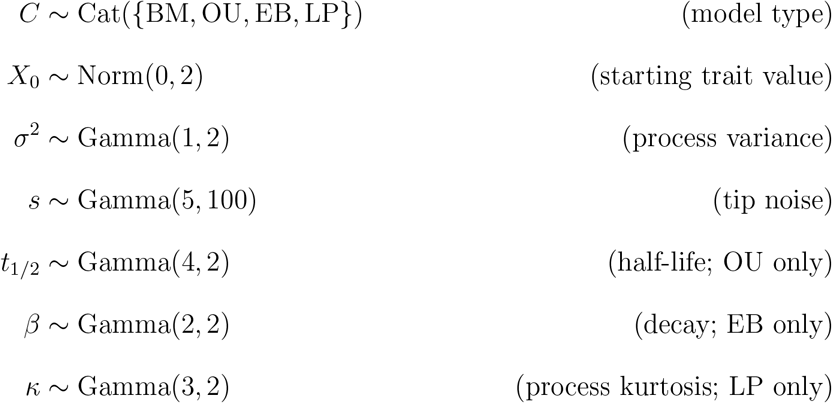

where the mean of Gamma(*α, β*) equals *α/β*. Process variance and kurtosis are the expected moments for trait change after one unit of time. For the OU process, process variance refers to the Brownian variance (i.e., not the stationary variance) and the optimal value is assumed to equal the ancestral value, *X*_0_. Phylogenetic half-life relates to the OU mean-reversion strength as *α* = ln(2)*/t*_1*/*2_ (Hansen 1997). Process variance and kurtosis are reparameterized into standard model parameters for LP (Landis and Schraiber 2017).

Note, these ranges of parameters were chosen so that datasets generated by more-complex models (OU, EB, LP) might be misinterpreted as originating from the simpler model (BM), thereby complicating model selection.

Trait datasets were simulated and fitted using the R package `pulsR` (Landis and Schraiber 2017). Maximum likelihood estimates were obtained from 5 independent Nelder-Mead optimization attempts. We then computed Akiake Information Criteria (AIC) scores for each model as *AIC*(*c*) = −2(*L*_*c*_ − *K*_*c*_), where *L*_*c*_ is the maximum log-likelihood and *K*_*c*_ is the number of free parameters for model *c*. We treated the model with the lowest *AIC*(*c*) value as the model selected by maximum likelihood (Akaike 1973).

To train `phyddle`, we simulated 200,000 datasets with a maximum tree width of 300. This analysis treated *C* as an estimation target, while ignoring other data-generating variables (e.g. *λ, µ, X*_0_, etc.) during training. We used CDV+S encoding with one row of numerical values for each species trait. We used a soft-max layer with a cross-entropy loss function to transform network output that model type *C* = *c* with probability *w*_*c*_ (Voznica et al. 2022). Interpreting *w*_*c*_ as proportional to the maximum (unlogged-) likelihood score for model *c*, we computed a penalized model selection score *SEL*(*c*) = −2(*ln*(*w*_*c*_) − *K*_*c*_). We then treated the model with the smallest value of *SEL*(*c*) as the model selected by `phyddle`.

We found that model selection performance between the two methods was similar: SEL with `phyddle` was correct 65.4% of the time, and AIC with `pulsR` was correct 65.8% of the time (Fig. 3). Both methods consistently select either the true model (diagonal) or the simpler nested model (BM, top row) in the majority of cases (Fig. 3A-B). Regardless of whether they selected the true model, the two methods selected the same model 71.7% of the time (Fig. 3C, diagonal). 21.7% of the time involved disagreement where one method selected the simplest model (BM). We note that `pulsR` requires the numerical integration of a recursively constructed characteristic function for model fitting, which grows prohibitively expensive for larger trees (see Landis and Schraiber 2017). `Phyddle` required approximately 10m to simulate and format data, 5m to train the network, and less than 5s to estimate parameters (15m total), whereas our parallelized maximum likelihood script needed about 5h to complete.

**Figure 3:**
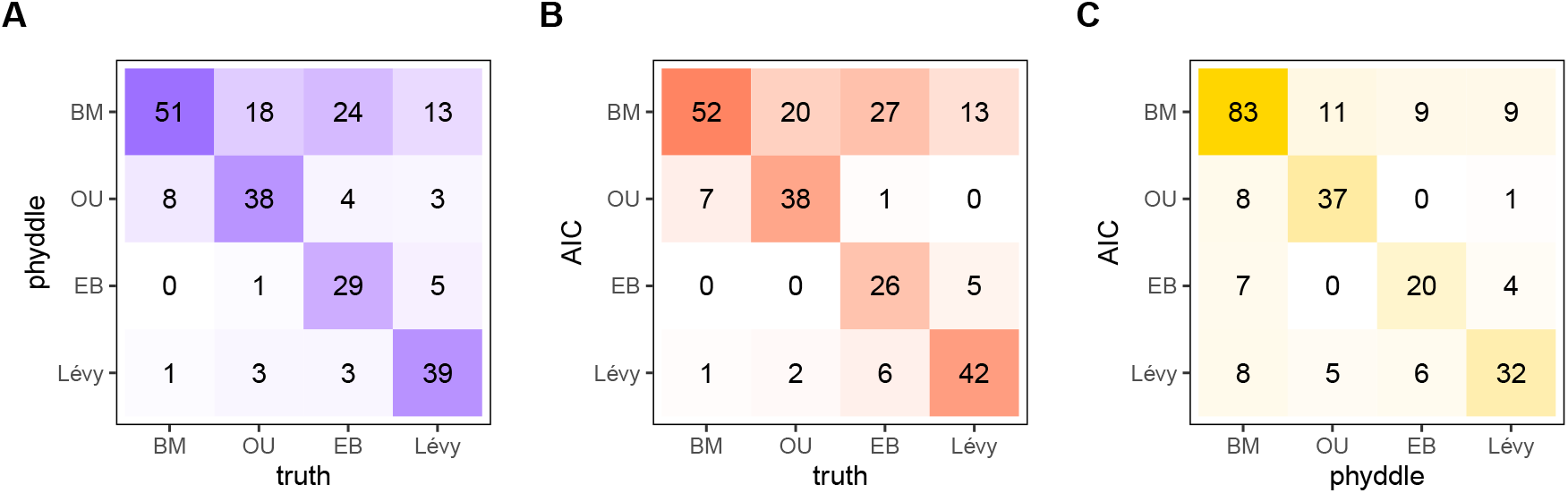
Comparison of model selection for AIC and `phyddle` estimates. Possible models are Brownian motion (BM), Ornstein-Uhlenbeck (OU), Early Burst (EB), and a Lévy process (LP; normal inverse-Gaussian). `phyddle` (A, blue) and AIC (B, red) reliably identify to true data-generating model, or the two methods agree (C, gold), when the frequencies along the diagonal are high. Off-diagonal entries represent model misidentification (A,B) or classification disagreement (C) for that column.

Finally, we showcase `phyddle`’s prediction interval capabilities using an evolutionary scenario that is not trivial for standard likelihood-based methods. We compared `phyddle` to the performance of likelihood-based inference methods under two Susceptible-Infected-Recovered (SIR) model scenarios. First, we simulated data for a phylogeographic model of pathogen infection that we refer to as a SIR + Migration or SIRM model (Riley 2007). We consider the case from (Thompson et al. 2024) where all pathogen sampling occurs during the exponential growth phase early in an outbreak, accomplished by rejecting simulations where more than 5% of susceptible individuals were infected. Only during the exponential growth phase, our SIRM model behaves as a serially sampled multitype birth-death model (Stadler and Bonhoeffer 2013; Kühnert et al. 2016), and is analyzed with a known and tractable likelihood function (Maddison et al. 2007; May and Meyer 2024). We used the test data and Bayesian estimates from (Thompson et al. 2024) to compare against our `phyddle` estimates.

We simulated outbreaks among five locations under the following conditions:

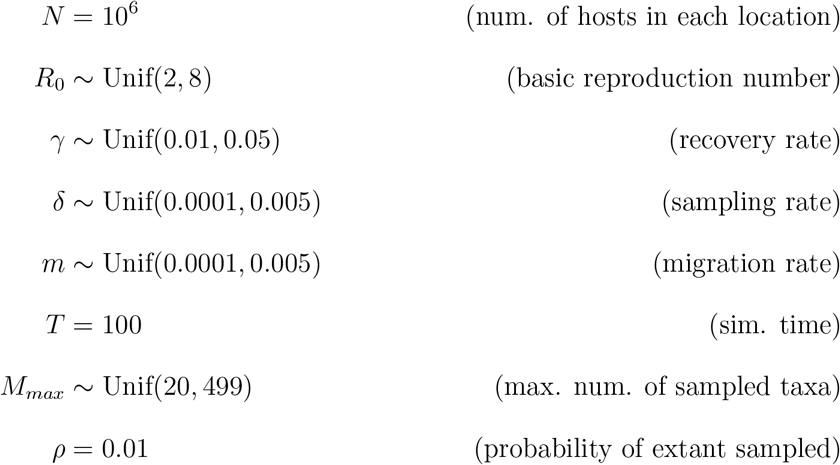

The per-host infection rate during the exponential-growth phase of an outbreak is *β* = *R*_0_(*γ* + *δ*). All simulations ran for *T* units of time, with each contagious individual at the end of the simulation (extant) having *ρ* probability of being sampled. The trees were downsampled to a maximum of *M*_*max*_ taxa.

In the second scenario, pathogens are sampled at a random time during the outbreak, both during and after the exponential growth phase; for this scenario we lack a simple likelihood-based inference strategy. We simulated training and test data under the same settings as above except we set:

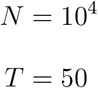

The smaller population size ensures many of the simulations extend beyond the exponential phase.

Note, the modeled scenarios are analogous, except that the data sampling window is expanded under the second scenario. We separately simulate data and train neural networks with `phyddle` for each scenario. We are not aware of an exact likelihood-based solution for the general SIRM model, i.e. where individuals can be sampled at any time during the outbreak and exact numbers of susceptible and infected individuals in location *i* at time *t* inform the local, instantaneous infection rate (but see Kühnert et al. 2016 and Müller et al. 2019). We emphasize that our comparison seeks to measure the accuracy of `phyddle` for a model with no simple likelihood-based counterpart. So, even if the unrestricted SIRM model had easily computable likelihoods, it would be trivial to engineer a more-realistic but less-tractable model (e.g. Moshiri et al. 2019; Shchur et al. 2022; Gavotte and Frutos 2022). As such, both our Bayesian analyses use the simpler, tractable, and time-constant SIRM model that assumes we only sample during the exponential growth phase.

We used RevBayes (HÖhna et al. 2016) with the TensorPhylo plugin (May and Meyer 2024) to estimate the joint posterior density of SIRM model parameters. We assumed the population size (*N*), recovery rate (*γ*) and extant sampling probability (*ρ*) were known empirically, while the reproduction number (*R*_0_), sampling rate (*δ*), and migration rate (*m*) were estimated. Prior distributions match the simulating distributions for all parameters. Markov chain Monte Carlo was run for 7,500 generations. A burnin period of 10% of generations was run before MCMC sampling and all model parameters had effective sample sizes of *>* 100.

We used `phyddle` to simulate 53,416 valid training examples for the exponential phase dataset and 96,555 for the all-phases dataset, as described above. Calibration datasets for performing conformal prediction by adjusting inner quantile estimates consisted of 6,677 for the exponential phase dataset and 37,136 for the all-phases dataset. Parameters *N* and *γ* were treated as known, whereas *R*_0_, *δ*, and *m* were inferred as free parameters. We used a CBLV+S tensor with one-hot encoding to represent phylogenetic and tip states across 5 locations.

Under the first scenario without model misspecification for the Bayesian approach, we see that both Bayesian and `phyddle` point estimates for the basic reproduction number are highly correlated with the true data-generating parameter values (Fig. 4A-B). In addition, lower and upper bounds for 95% highest posterior density credible intervals and conformalized prediction intervals are tightly correlated (Fig. 4C-D).

**Figure 4:**
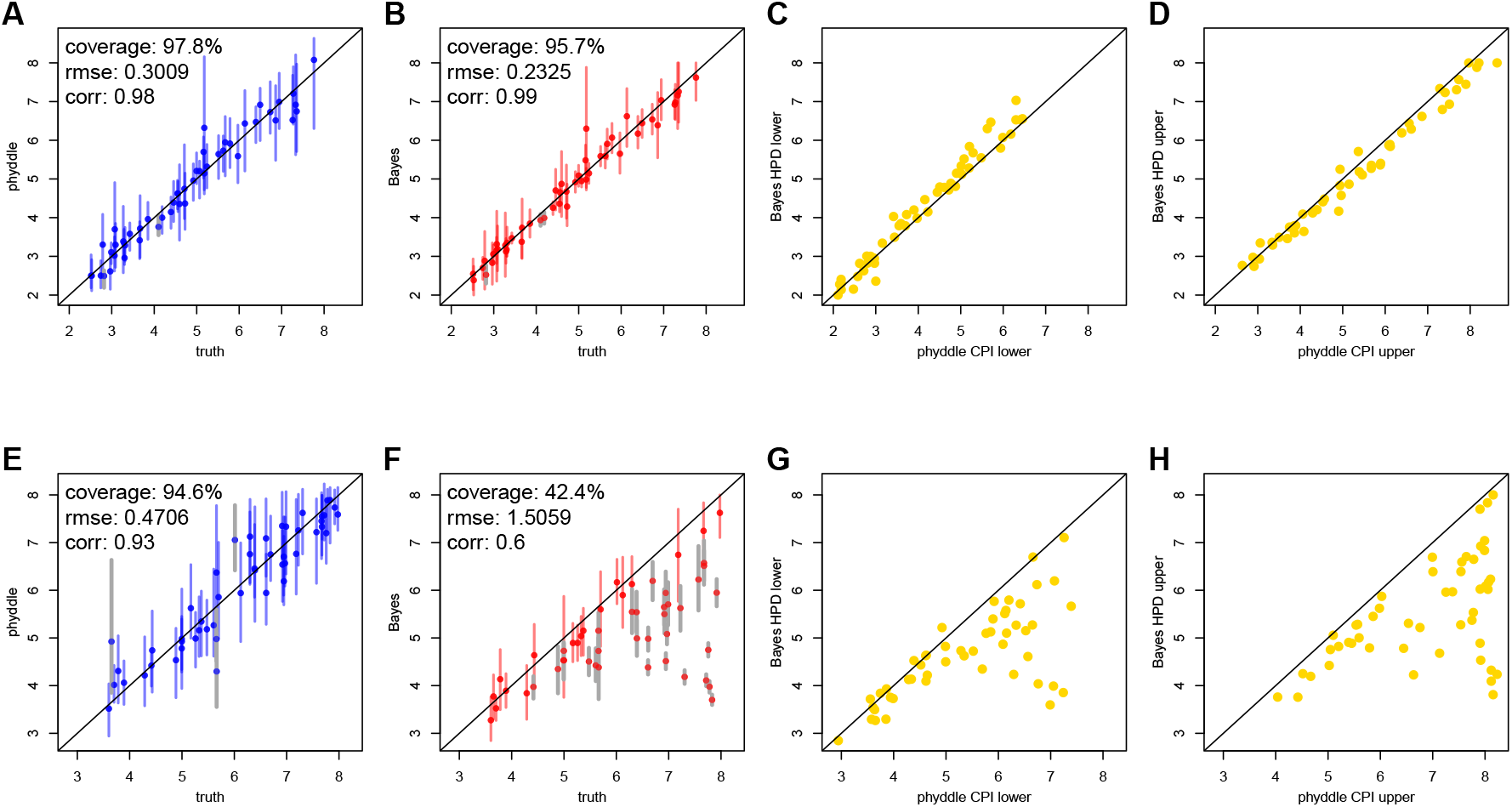
Comparison of Bayesian and `phyddle` estimates for the basic reproduction number, *R*_0_, when all pathogens are sampled during the exponential growth phase of an outbreak (A-D) or sampled at any time during an outbreak (E - H). True parameter values are plotted against `phyddle` (blue; A and E) and Bayesian (red; B and F) point estimates. Estimated support interval bounds (gold; C, D, G, and H) for `phyddle` and Bayesian methods are also plotted against each other. Any point that falls on a slope-1 intercept-0 line has perfectly matching *x* and *y* values. Data displayed is a random subsample of 50 values (roughly 50%). Intervals shown are 95% CPI (conformalized prediction interval) or HPD (highest posterior density). Bayesian estimates and test data for comparison of exponential phase data (A-D) are from (Thompson et al. 2024).

Under the full outbreak scenario, `phyddle` accurately estimates the true data-generating parameters (Fig. 4E), but the Bayesian posterior point estimates and support interval bounds no longer agree with `phyddle` (Fig. 4F-H). That is, as the outbreak progresses, the number of new hosts that are still susceptible to infection decreases, so the infection rate slows, hence the consistent underestimation for the reproductive number when the model assumes unbounded, exponential growth in the number of infections.

Similar results for the sampling rate, *δ*, and migration rate, *m*, are provided in the Supplement. This behavior is expected because, although the neural networks trained by `phyddle` conform to all model assumptions encoded into the simulator, the simulator violates the simplifying model assumptions required to obtain a tractable likelihood for our Bayesian analysis.

For the full analysis, `phyddle` required approximately 10h to simulate and format data, 20m to train the network, and less than 10s to estimate parameters (10.5h total). Our Bayesian analyses took about 2h to complete, when run in parallel; we did not have a tractable likelihood for the model under the full outbreak scenario.

Although this experiment only demonstrates how inference methods behave when models are correctly versus incorrectly specified for two epidemiological scenarios, the broader need to mitigate model misspecification and potential solution offered by likelihood-free methods, such as `phyddle`, are more general.

## Summary

We expect that `phyddle` has many possible uses for researchers, both green and seasoned, applied and theoretical. We name a few ideal use cases. First, `phyddle` allows biologists to design and fit phylogenetic models with no known, tractable inference methods, which increases community-level access to newer models worthy of attention. Second, `phyddle` enables users to rapidly test whether a dataset contains any signal under standard models that have likelihood-based inference methods. A one-day exploratory analysis could inform a biologist whether they should spend a year collecting data and/or developing new methods. Third, `phyddle` scales well for repetitive estimation tasks, making it ideal for analyses involving many clades, many genes, and/or assessing model sensitivity to phylogenetic uncertainty. Fourth, `phyddle` can be used to provide baseline estimates when developing new likelihood-based approach with no existing points of comparison to benchmark performance. Lastly, because deep learning in phylogenetics is currently underexplored, the customizability of `phyddle` can help researchers to assess how various modeling, network architecture, and training conditions influence the effectiveness of deep learning approaches, e.g. through simulation experiments.

It remains unknown how well deep learning will perform in uncharted regions of model complexity. Results from Thompson et al. (2024) suggest that deep learning approaches, similar to those employed by `phyddle`, can match the performance of likelihood-based approaches when evaluating inference models with tractable likelihoods. This implies deep learning might perform comparably well for models that have no known likelihood functions. With this in mind, `phyddle` is designed to help researchers rapidly, safely, and systematically apply deep learning approaches to phylogenetic model exploration and data analysis.

## Software

`phyddle` is an open source project that is written in Python. Code for `phyddle` is hosted at https://github.com/mlandis/phyddle. The software depends on a large number of scientific computing libraries, including: `pytorch` (Paszke et al. 2019), `dendropy` (Sukumaran and Holder 2010), `numpy` (Harris et al. 2020), `scipy` (Virtanen et al. 2020), `scikit-learn` (Pedregosa et al. 2011), `pandas` (pandas development team 2020), `h5py` (Collette 2013), and `matplotlib` (Hunter 2007). Documentation and tutorials for `phyddle` are hosted at https://phyddle.org. Saved workspace projects for the three examples in this manuscript are hosted at https://github.com/mlandis/phyddle_ms.

## Supplementary Information

Supplementary Information is hosted on Dryad: https://datadryad.org/stash/share/k7d0ndDkKiVx-fK2B81WaJ5Mk_vn7WUKv2nynfbO8Ug.

## Acknowledgements

We are grateful to Albert Soewongsono, Raymond Castillo, Sean McHugh, Sarah Swiston, Fábio Mendes, Sigournie Brock, Tracy Heath, and Erik Scully for feedback and help testing software. Insightful comments from four anonymous reviewers, the associate editor, and the editor-in-chief also helped improve the quality of the article.

