## Supplementary Information for "phyddle: software for exploring phylogenetic models with deep learning"

Version dated: February 27, 2025

RH: phylogenetic model exploration with phyddle

### Supplementary Information: phyddle: simulation-trained deep learning predictions for phylogenetic models

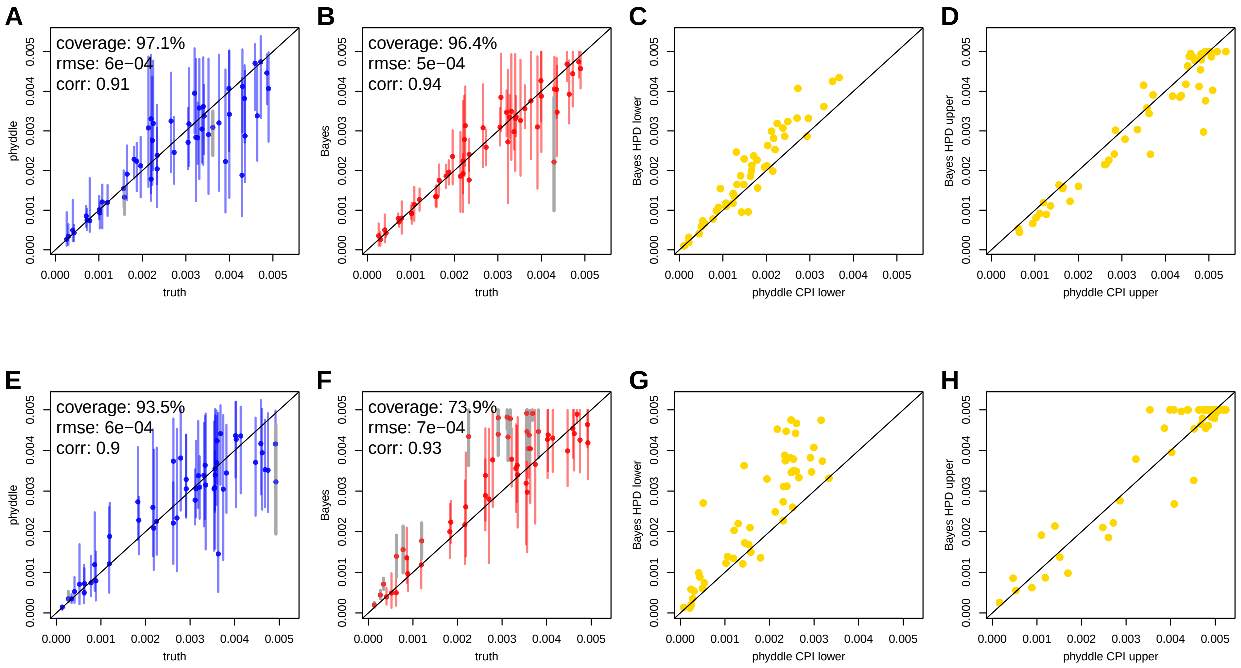


Figure S1: Comparison of Bayesian and `phyddle` estimates for the sampling rate, $\delta$, when all pathogens are sampled during the exponential growth phase of an outbreak (A-D) or sampled at any time during an outbreak (E - H). True parameter values are plotted against `phyddle` (blue; A and E) and Bayesian (red; B and F) point estimates. Estimated support interval bounds (gold; C, D, G, and H) for `phyddle` and Bayesian methods are also plotted against each other. Any point that falls on a slope-1 intercept-0 line has perfectly matching $x$ and $y$ values. Data displayed is a random subsample of 50 values (roughly 50%). Intervals shown are 95% CPI (conformalized prediction interval) or HPD (highest posterior density). Bayesian estimates and test data for comparison of exponential phase data (A-D) are from (Thompson et al. 2024). See main text for analysis details (Landis and Thompson 2024).

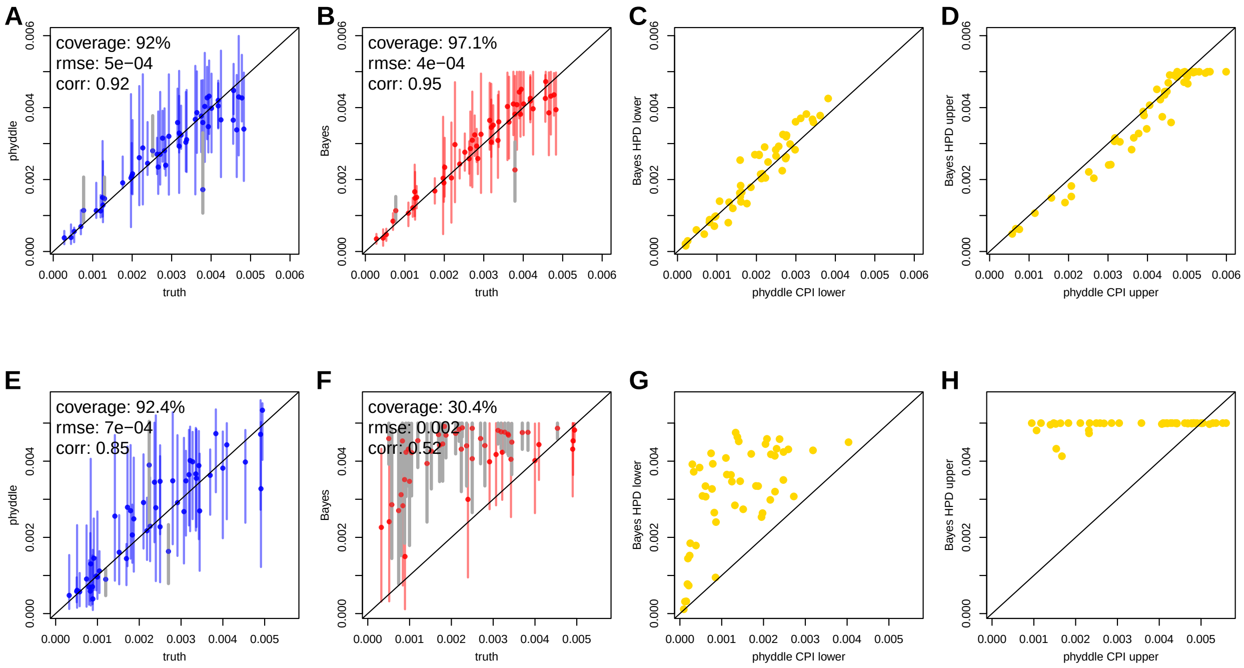


Figure S2: Comparison of Bayesian and `phyddle` estimates for the migration rate, $m$, when all pathogens are sampled during the exponential growth phase of an outbreak (A-D) or sampled at any time during an outbreak (E - H). True parameter values are plotted against `phyddle` (blue; A and E) and Bayesian (red; B and F) point estimates. Estimated support interval bounds (gold; C, D, G, and H) for `phyddle` and Bayesian methods are also plotted against each other. Any point that falls on a slope-1 intercept-0 line has perfectly matching $x$ and $y$ values. Data displayed is a random subsample of 50 values (roughly 50%). Intervals shown are 95% CPI (conformalized prediction interval) or HPD (highest posterior density). Bayesian estimates and test data for comparison of exponential phase data (A-D) are from (Thompson et al. 2024). See main text for analysis details (Landis and Thompson 2024).

*
